# Bioengineered extracellular vesicles mitigate neuroinflammation by neutralizing pneumolysin and delaying disease onset in experimental pneumococcal meningitis

**DOI:** 10.64898/2026.01.07.698151

**Authors:** Kristine Farmen, Miguel Tofiño-Vian, Doste R. Mamand, Xiuming Liang, Houze Zhou, Vicky W.Q. Hou, Samir El Andaloussi, Oscar P. B. Wiklander, Federico Iovino

## Abstract

Bacterial meningitis is a life-threatening neurological disorder frequently caused by a *Streptococcus pneumoniae* (the pneumococcus) infection of the brain. Standard treatment consists of antibiotics to eliminate bacteria and dexamethasone to reduce inflammation. Despite this, mortality reaches 20% in treated individuals, and half of the survivors suffer long-term neurological sequelae. This is largely due to the poor capacity of antibiotics to reach the brain and the lack of antimicrobial treatment capable of neutralizing the pneumococcal toxin pneumolysin (Ply). To address these limitations, we isolated extracellular vesicles (EVs) derived from human HEK293T cells and evaluated their therapeutic potential in pneumococcal meningitis. Alongside wild-type EVs (WT.EVs), we bioengineered EVs to express RVG peptides (RVG.EV) for targeting neuronal acetylcholine receptors, signal incompetent IL-6 signal transducer (IL-6ST) decoy receptors (IL-6.EV) to block the pro-inflammatory signalling of IL-6, or EVs expressing both RVG peptides and IL-6ST (DB.EV). *In vitro,* all EVs reduced pneumococcal adhesion to neurons and mitigated cytotoxicity by binding and sequestering Ply. In a bacteremia-derived pneumococcal meningitis model, EV treatment significantly increased the survival of the mice without affecting bacterial load in the brain or the periphery. Among all groups, RVG.EV treatment was most effective in reducing pro-inflammatory cytokine release in the periphery and brain. These findings highlight the therapeutic potential of bioengineered EVs, particularly RVG peptides expressing EVs, as an adjunctive treatment for pneumococcal meningitis thanks to their (i) sequestration and neutralization of Ply, (ii) increased blood-brain barrier crossing, and (iii) dampening of inflammation.

## 1. Introduction

*Streptococcus pneumoniae* (the pneumococcus) infections are responsible for one million deaths of children worldwide annually [1]. The most serious manifestation of the disease is invasive pneumococcal disease, where the bacterium enters sterile body sites such as the blood and brain, causing bacteremia and meningitis, respectively [2]. *S. pneumoniae* is a major cause of bacterial meningitis worldwide and is associated with a worse patient prognosis [3, 4]. Survival is dependent on rapid access to treatment; however, even among treated individuals, a mortality rate of 20% is observed, and about 50% of survivors experience permanent neurological sequelae due to neuronal damage inflicted by bacteria and bacterial toxic components [5, 6]. This poor prognosis is driven by challenges with antibiotic treatment, including antibiotic resistance, the low ability of antibiotics to cross the blood-brain barrier (BBB), which leaves the brain untreated for a prolonged period, and exacerbated neuroinflammation due to bacterial lysis by antibiotics [7, 8].

*S. pneumoniae* activates inflammatory cascades, leading to the release of pro-inflammatory cytokines and chemokines [9]. Elevated TNF-α and IL-6 levels in the cerebrospinal fluid (CSF) correlate with disease severity in patients, and the release of chemokines increases the infiltration of peripheral monocytes and neutrophils to aid microglia in the clearance of the infection; however, the clearance is inefficient, and a prolonged inflammatory state occurs [10, 11]. This is detrimental as inflammatory components such as matrix metalloproteinases and reactive oxidative species are toxic to cells [12]. The upheld neuroinflammation is associated with increased intracranial pressure, alteration in blood flow, herniation, and infarction [5, 13]. Dexamethasone is used to mitigate neuroinflammation and has been shown to improve patient outcomes [14].

In addition to the host-mediated damage, *S. pneumoniae* causes cellular damage by releasing pneumolysin (Ply) [15]. Ply is a potent stimulator of pro-inflammatory cytokines through interaction with Toll-like receptor (TLR) 2 and 4 on immune cells [16]. The toxin also aids *S. pneumoniae* evasion from microglial cells by reducing their motility [17]. Ply is a cholesterol-dependent cytolysin, and the interaction with membrane cholesterol generates a pore in the cell membrane, causing cell lysis [18–20]. In addition, Ply facilitates the binding of *S. pneumoniae* to neuronal cells, disrupting the cytoskeleton, leading to cell death [21, 22]. Further, it contributes to hearing loss and cortical damage in experimental pneumococcal meningitis [23, 24]. Ply also disrupts the ciliary beating in ependymal cells and causes cytoskeleton remodeling in astrocytes, potentially affecting the CSF flow in the brain and contributing to brain edema [25, 26]. Importantly, increased Ply concentrations in the brain correlate with increased risk of death in patients [27].

Since the cellular damage observed in pneumococcal meningitis is a consequence of the neuroinflammatory process and direct damage by *S. pneumoniae*, novel therapies must effectively and quickly reach the brain, reduce *S. pneumoniae* cytotoxicity towards cells by blocking direct interaction, neutralize Ply, and reduce inflammation. Extracellular vesicles (EVs) have the potential to fulfil all of these requirements thanks to their intrinsic beneficial properties; they exhibit low immunogenicity and are rich in cholesterol and thus putatively bind Ply [28]. Furthermore, EVs cross cellular barriers, including the BBB, the latter is increased during systemic inflammation [29, 30]. Importantly, they can be bioengineered to express receptors and peptides [31].

We focused on two such strategies: 1) Bioengineering of EVs with rabies viral glycoprotein (RVG) peptides that bind to the nicotinic acetylcholine (nACh) receptors widely expressed by neurons [32, 33]. This expression of RVG was reported to cause a two-fold increase in brain invasion compared to wild-type EVs [32, 34]. Thus, we hypothesized that EVs expressing RVG peptides could act as a molecular barrier between neurons and *S. pneumoniae,* reducing the cellular cytotoxicity during infection. 2) Bioengineering EVs expressing signal incompetent IL-6 signal transducer (IL-6ST) decoy receptors that block the pro-inflammatory trans-signalling pathway of IL-6 [35]. 3) Generating double-coated EVs expressing both RVG peptide and IL-6ST, hypothesizing that this inhibition would lower both neurotoxicity and inflammation. We also speculated that the EVs could, due to their membrane cholesterol, bind Ply and thus reduce cellular cytotoxicity. Therefore, we assessed the therapeutic potential of wild-type EVs (WT.EV) or bioengineered EVs expressing either RVG peptides (RVG.EV) or signal incompetent IL-6ST decoy receptors (IL-6.EV) or both RVG.IL-6EVs (DB.EV) from HEK293T cells in an experimental pneumococcal meningitis model.

*In vitro,* we demonstrated that all EVs sequestered Ply and reduced *S. pneumoniae*-induced neuronal cytotoxicity during infection. Further, by inhibiting trans-signalling of IL-6, the release of pro-inflammatory cytokines was reduced. In a bacteremia-derived meningitis mouse model, all EV types had a protective effect and delayed disease onset. Interestingly, the RVG.EV treatment outperformed the rest of the treatment groups and reduced both microglial activation and release of pro-inflammatory cytokines in the brain and periphery. Ultimately, we demonstrated that treatment with bioengineered EVs expressing RVG peptides in pneumococcal meningitis was effective and holds vast potential as an adjuvant treatment in severe pneumococcal infections.

## 2. Materials and methods

### 2.1 EV production and isolation

In total four different EV types were generated: wild-type EVs (WT.EV), the single expressing EVs RVG.EV and IL-6.EV, containing the RVG peptides and signal incompetent IL-6ST decoy receptors respectively, and finally the double-expressing EVs containing both RVG peptides and the signal incompetent IL-6ST decoy receptors (DB.EVs). The EVs were generated as previously described [34–36]. Briefly, EVs were generated through polyethylenimine (PEI)-mediated transient transfection of HEK293T cells. Cells were initially seeded in 15-cm dishes at a density of 10 × 10⁶ cells per dish using complete DMEM (10 % FBS and 1% penicillin-streptomycin-antimycotic) (Gibco,). After 48 hours, cells were transfected with the appropriate plasmids. Six hours post-transfection, the culture medium was replaced with Opti-MEM (Gibco) supplemented with 1% penicillin-streptomycin-antimycotic. After an additional 48 hours, the conditioned medium (CM) was harvested and purified by sequential centrifugation at 700 × g for 5 min and 2,000 × g for 10 min. The resulting supernatant was passed through a 0.22 µm filter to remove cellular debris and large particles.

EVs were isolated from a filtered conditioned medium using tangential flow filtration (TFF) using a MicroKross system (20 cm², Spectrum Labs) equipped with a 300 kDa molecular weight cutoff membrane, allowing for selective retention and concentration of vesicle-sized particles. The retentate was further concentrated using Amicon Ultra-15 centrifugal filters (100 kDa cutoff, Millipore) by centrifugation at 4,000 × g at 4°C. Spin durations ranged from 30 min to several hours, depending on the sample volume and EV content. The resulting EV concentrate was transferred into maximum recovery 1.5ml microtubes (Axygene,) and analyzed by Nanoparticle Tracking Analysis to determine particle concentration and size distribution.

### 2.2 EV characterization

*Nanoparticle tracking analysis (NTA).* NTA measurements were conducted with the Zetaview system (Particle Metrix). All samples were diluted in 0,22µm filtered PBS to achieve a final volume of 1ml. The ideal measurement concentrations were determined through pre-testing to achieve a particle count of 140–200 particles per frame. The manufacturer’s default software settings for EVs were selected accordingly. In brief, the settings included a sensitivity of 80 and a shutter speed of 100. Two cycles were performed for each measurement, scanning 11 cell positions and capturing 60 frames per position (with the video setting set to high). The specific settings for the measurements were as follows: Focus was set to autofocus; camera sensitivity was set to 80.0; shutter speed remained at 100; scattering intensity was adjusted to 4.0; and the cell temperature was maintained at 24°C. After video capture, the data were analyzed using the built-in ZetaView Software version 8.02.31.

*Western blotting:* The intravesicular protein levels of TG101, surface RVG and Gp130 (IL-6ST) in engineered EVs were determined using western blot analysis, as previously described [37]. Calnexin was used as a marker for cell lysates and to assess the purity of EVs. In brief, 2 × 10^10^ EVs and 2 × 10^6^ HEK293T cells were lysed in 100µl of radioimmunoprecipitation buffer (RIPA; BioRad, Hercules). The samples were incubated on ice for 30 min, vortexed for 10 seconds every 5 min. Afterward, the EVs and cell lysates were mixed with 8µl of loading buffer (10% glycerol, 8% sodium dodecyl sulfate, 0.5 M dithiothreitol, and 0.4 M sodium carbonate). This mixture was incubated at 65°C for 5 min, loaded onto a NuPAGE+ (Invitrogen, Novex 4-12% Bis-Tris gel), and run at 120 V for 1.5 hours. The proteins were transferred to an iBlot membrane (iBlot 2 Transfer Stacks; Invitrogen) using the iBlot 2 Dry Blotting System for 7 min. The membrane was then incubated with blocking buffer (Odyssey Blocking Buffer; LI-COR Biosciences) at room temperature (RT) for 1 hour and subsequently incubated overnight at 4°C with freshly prepared primary antibodies: anti-RVG (1:1000, Invitrogen, Cat#:PA5-117507), anti-TSG101 (1:500, Invitrogen, Cat#:MA1-23296) anti-calnexin (1:000, Invitrogen Cat#: MCA497G), and anti-Gp130 (IL-6ST) (1:1000, Invitrogen, Cat#14-5982-82). The membranes were washed four times with TBS-T 0.1% for 5 min each time on a shaker. Following the washes, the membranes were incubated for 60 min with a 1:10000 diluted secondary antibody (IRDye® 800CW Goat anti-Mouse IgG, Goat anti-Rabbit IgG and Donkey anti-Goat IgG, LI-COR Biosciences). The membranes were washed three times with PBS, and the results were visualized using an infrared imaging system (LI-COR Odyssey CLx).

The protein levels of Ply, CD63 and CD81 used to assess EV-Ply binding and EV brain tissue infiltration was performed as previously described [22]. Briefly, samples were resuspended in LDS sample buffer (1:4, Invitrogen) and boiled at 95°C for 10 min and loaded onto NuPage+ (Invitrogen, Novex 4-12% Bis-Tris gel). Electroblotting was performed using the Mini Gel Tank (Thermo Fisher Scientific). Samples were blocked with PBS-T (T=0,1% Tween) supplemented with 5% skim-milk powder (Merck) for 1 hour at RT; and incubated with primary antibodies: Ply (1:1000 Abcam), CD63 (1:1000, Abcam), CD81 (1:1000, Invitrogen) overnight. Membranes were washed in PBS-T and incubated with secondary antibodies (1:5000, Goat anti-Rabbit IgG (H+L), Invitrogen) for 1 hour. Both primary and secondary antibodies were diluted in PBS-T supplemented with 2.5% skim-milk powder (Merck). Protein bands were detected by incubating membranes with ECL Prime Western Blotting Detection Reagents (Cytiva) and imaged using ImageQuant LAS 4000 (GE Healthcare). For Coomassie staining, gels were incubated with InstantBlue Coomassie Protein Stain (Abcam) overnight and imaged. Band intensities were measured using the analyze gels plug-in and the area under the curve (AUC) using ImageJ (Fiji). For ultracentrifugation experiments, the total Ply AUC was divided by the total AUC for Coomassie staining, for qEV Ply experiment the Ply AUC was divided by the CD63 AUC. The sequestering capacity of the EVs was calculated as Pellet Ply/ Total Ply (Pellet + supernatant fraction).

*Single-vesicle imaging flow cytometry for characterization of EV surface markers.* Characterization of EV Surface markers using flow cytometry was done as previously described [33]. Shortly, purified Evs were diluted to a concentration of 1 × 10^10^ particles/ml using Dulbecco’s Phosphate-Buffered Saline. A total of 2.5 × 10^8^ EVs in 25 µl were stained with either 8 nM of phycoerythrin (PE), fluorescein isothiocyanate (FITC), or Allophycocyanin (APC) conjugated antibodies targeting the surface markers or proteins of interest, RVG-APC (Novus Biologicals), and Gp130-APC (IL-6) (eBioscience), CD9-PE (Miltenyi Biotec), CD63-PE (Miltenyi Biotec), or CD81-FITC (Miltenyi Biotec), which are known to be enriched in EVs. The EVs were then incubated overnight at room temperature in the dark within a sealed 96-well plate (SARSTEDT AG & Co. KG). After incubation, the samples were diluted in a 1% human albumin and trehalose (PBS-HAT) buffer solution to achieve a final concentration of 1 × 10^7^ EVs/ml in a total volume of 100 µl [34]. Single-vesicle imaging was performed using an Amnis Cellstream (Luminex) and the results examined in FlowJo (v. 10.8; FlowJo LLC).

*Transmission electron microscopy.* 3μl of each EV sample was applied to glow-discharged, carbon-coated, formvar-stabilized 400 mesh copper grids (easiGlow™, Ted Pella) and incubated for approximately 30 sec. The excess sample was blotted off, followed by a MilliQ water wash. Grids were negatively stained with 1% ammonium molybdate and imaged using a HT7800-Xarosa transmission electron microscope (Hitachi High-Technologies) operated at 80 kV, equipped with a 4 MP Veleta CCD camera (Olympus Soft Imaging Solutions GmbH).

### 2.3 Cultivation of Streptococcus pneumoniae

*Streptococcus pneumoniae* TIGR4 (serotype 4) (T4) was grown to exponential phase (OD of 0.3) in Todd-Hewitt broth (THY, Substrat, KI) and stored in THY solution supplemented with 20% glycerol (Substrat, KI) at −80°C. Before *in vitro* experiments, the bacterial aliquots were thawed, pelleted, and washed in PBS to remove any trace of glycerol, and resuspended in cell culture media. The day before the *in vivo* experiments, the aliquots were thawed, pelleted, resuspended in PBS, and frozen overnight at −20°C.

### 2.4 Cell culture

Human SH-SY5Y neuroblastoma cells were cultured as previously described [21]. Briefly, cells were seeded in 6-well plates at a density of 300.000 cells/well, or 100.000 cells/well in 12-well plates, differentiated for 12 days to obtain mature neuronal cells using growth media made of equal parts EMEM (ATCC) and F12 (Gibco), 5% heat inactivated fetal bovine serum (FBS) (Gibco), 1% penicillin/streptomycin (Gibco), and 10μM retinoic acid (RA) (Biotechne). The BV-2 mouse microglial cell line (Cytion, Eppelheim,) was plated at a density of 300.000 cells/well in 6-well plates in DMEM medium (Gibco) supplemented with 2.5% heat-inactivated FBS (Gibco) and 1% penicillin/streptomycin (Gibco). All cell lines were incubated at 37°C with 5% CO_2_. Media was changed every 2^nd^ or 3^rd^ day. One day before infection experiments, the cells were changed to their respective growth media without antibiotics.

### 2.5 *In vitro* bacterial adhesion and cytotoxicity assays

*In vitro* assays for bacterial adhesion and cytotoxicity were performed as previously described [22, 38]. Shortly, on the day of the experiment, the cell medium was discarded, and cells were washed in pre-warmed PBS and changed to pre-warmed cell medium without RA for SH-SY5Y and antibiotics for both cell lines. The cells were incubated for 1 hour at 37°C in 5% CO_2_ to acclimatize. Cells were infected with a Multiplicity of Infection (MOI)=10 for neuronal cells and 20 for BV-2 cells and treated with 1 x10^11^ p/ml of EVs or equal volumes of PBS (control) at the same time as infection (EV:cell ratio =10^6^:1) or added 30 min post infection (EV:cell ratio = 10000:1). In assays with antibiotic treatment, a dosage of 25μg/ml Ceftriaxone (Navamedic) was added 30 min post-infection. Infected cells were incubated at 37°C and 5% CO_2_ for 2 hours, subsequently, the supernatants were collected (non-adhered bacteria), and cells were washed with PBS to eliminate unbound bacteria. Next, cells were treated with 1ml of trypsin for 15 min, and the cell suspension was collected (adhered bacteria). Serial dilutions were performed of the non-adhered and adhered fractions and plated on blood agar plates overnight. The following day the Colony Forming Units (CFUs) were determined. The percentages of adhered bacteria were determined as the (adhered bacteria)/(non-adhered + adhered bacteria) x 100%.

For the cytotoxicity assays, the CyQUANT LDH Cytotoxicity Assay Kit (Invitrogen, Cat#C20300 and C20301) was used to measure the released lactase dehydrogenase (LDH), as an assessment of cytotoxicity. Infection was performed as previously described, while for assessment of Ply sequestering and cytotoxicity, SH-SY5Y neuronal cells were treated with 2.0µg/ml recombinant Ply (Protein Production Platform, Nanyang Technological University, Singapore), in combination with 1 x 10^9^ p/ml of EV groups, controls were treated with PBS for 45 min (EV:Cell ratio = 10000:1). After infection/treatment, 50μl from the supernatant of each sample was transferred to a 96-well flat-bottom plate, in technical triplicates. Untreated and uninfected cells were used as negative controls; the positive controls were cells treated with lysate buffer (Invitrogen). Absorbance was measured with Multiskan FC microplate reader (Thermo Fisher Scientific) at 450 nm and 620 nm; to determine LDH activity, 620 nm absorbance value (background from the instrument) was subtracted to 450 nm absorbance value.

### 2.6 Scanning electron microscopy (SEM)

SH-SY5Y-differentiated neuronal cells were grown on glass coverslips (Fisher Scientific) and infected with T4 at MOI=50 and treated with 2,3 x 10^11^ p/ml of DB.EVs for 30 minutes at 37°C with 5% CO_2_; untreated cells were used as a negative control (EV: Cell ratio =10^6^:1). The cells were fixed by immersion in 2.5 % glutaraldehyde (Ladd Research) in 0.1M phosphate buffer, pH 7.4. The coverslips were rinsed in phosphate buffer pH 7.4 followed by MilliQ water, followed by ethanol dehydration and critical point drying using carbon dioxide (Leica EM CPD 030). The coverslips were mounted on alumina SEM pins using carbon conductive tabs (Ted Pella) and sputter coated with a 10 nm layer of platina (Quorum Q150T ES). The images were acquired using an Ultra 55 field emission SEM (Zeiss) at 3 kV and the SE2 detector.

### 2.7 EV-Ply binding assay

To assess the sequestering capacity of EVs, 2µg/ml of Ply (Protein Production Platform, Nanyang Technological University, Singapore) was incubated with 2.2 x 10^9^ p/ml EVs for 30 min on ice in Ultra Clear Tubes (Beckmann Coulter) and placed in a precooled ultracentrifuge (Beckmann Coulter). The samples were centrifuged for 70 min at 100 000 x g at 4°C. The supernatant and pellet were collected separately, and BCA was performed to determine protein concentration prior to western blot analysis (described section 2.2).

### 2.8 Size exclusion chromatography (SEC) for EV-Ply binding

70 nm qEV original columns (500 µl; IZON Science Ltd) were equilibrated according to the manufacturer’s instructions. A mixture of 2.2 x 10^11^ p/ml of respective EV type and 2µg/ml of Ply (Protein Production Platform, Nanyang Technological University, Singapore), pre-mixed for 15 min RT, was loaded onto each column. The EV-enriched fraction (4^th^ and 5^th^ ml eluted fractions) was collected, concentrated to a final volume of 200–300µl using 2 ml 10 kDa molecular weight cut-off spin filters (Amicon Ultra; Millipore) and centrifuged at 4,000 × g at 4°C. The concentrated EVs were analyzed with NTA and the Ply concentration in each elute was determined by western blot analysis (described in section 2.2)

### 2.9 Hemolytic activity

For all hemolytic assays pooled human red blood cells (Medix biochemical) were washed by diluting 1:10 in PBS (Substrat) and centrifuged 500 x g 10 min at RT. The pellet was resuspended 1:100 in PBS. Hemolysis was induced by adding recombinant Ply (Protein Production Platform, Nanyang Technological University, Singapore) in the presence of EVs (concentrations are stated in figure legends). PBS alone was used as a negative control, and 0.03% Triton X-100 (Merck) as a positive control. The solutions were placed on a shaking incubator at 37°C for 45min, and non-lysed red blood cells were pelleted by centrifugation 500 x g 10 min at RT. The supernatant was collected, and hemoglobin release was determined by absorbance measurement with Multiskan FC microplate reader (Thermo Fisher Scientific) at 450 nm. The percentage of hemolysis was calculated as follows: A_450_ sample-OD A_450_negative control/ (A_450_positive control- A_450_negative control). All experiments were conducted with 3 technical replicates.

### 2.10 Animal experiments

Five- to six-week-old male C57BL/6J mice (JAX™, Charles River) were housed in a 13:11 hour light/dark period with *ad libitum* access to water and food in accordance with Swedish legislation (Svenska jordbruksverket, ethical permit number 18965-2021). We utilized a bacteremia-derived meningitis model as previously described [39, 40]. For the first pilot study: Each mouse received an intravenous injection of 1 x 10^8^ CFU of *S. pneumoniae* in PBS solution through the tail vein, control animals received PBS. 1 hour later, mice received treatment with 0.5-2 x10^11^ particles/mouse WT.EVs through an I.V injection. Control animals received PBS. The mice were assessed for clinical symptoms every 3 hours and scored according to Karolinska Institutet’s ‘Assessment of health conditions of small rodents and rabbits when illness is suspected’ template. The experimental endpoint was set 8 hours post-T4 injection, to assess the effect of the EVs on bacterial growth *in vivo*. For the other animal experiments, the infection and treatment time points were the same, however, the treatment dosage was 2 x10^11^ particles/ mouse for all EVs and the experimental endpoint was when the animal reached a score of 0.4-0.5, at which time the animals displayed both neurological and systemic symptoms of the disease. An overview of the different animal studies, the purpose, type of EV treatment, and downstream analysis is depicted in **Supplementary Table 1**.

At the time of sacrifice, blood was collected for assessment of bacterial load and/or cytokine expression. The mice were perfused with ice-cold PBS through the left ventricle, and the brain was collected and immediately dissected into two hemispheres. The left hemisphere was placed in ice-cold 4% PFA (Histolab) for immunohistology downstream analysis, while the right hemisphere was placed in PBS for preparation of brain homogenates. The spleen was collected and placed in ice-cold PBS. Spleen and brain tissue were homogenized through a 100µm cell strainer and frozen at −80°C with 1% protease inhibitor cocktail added (Fisher Scientific). Serial dilutions of tissues and blood were plated on blood agar plates (Substrat) for the determination of CFU numbers and incubated overnight at 37°C with 5% CO_2_

### 2.11 Immunostaining and confocal microscopy

The left hemisphere of the brain was post-fixed overnight at 4°C, placed in a 30% sucrose (Merck) solution for a minimum of 48 h at 4 °C, and subsequently cut into coronal sections of 30μm on a microtome (Leica SM2000). The sections were frozen in a cryoprotective solution (30% sucrose and 2% DMSO (Merck) in PBS) and stored at −20°C until further processing. Before staining, the sections were washed in PBS, blocked with 5% goat serum (Gibco), and permeabilized with 0.3% Triton-X-100 (Merck) for 1 hour at RT and stained with primary antibodies overnight at 4 °C. After washing with PBS, the sections were incubated with secondary antibodies for 2 hours at RT. The primary antibodies used in this study were: Iba1 (1:1000 Abcam, Cat#:MA5-36257), GFAP (1:250, Cell Signaling Technologies, Cat#:3670), and NeuN (1:250, Proteintech, Cat#:66836-1-Ig). Secondary antibodies used were Alexa Fluor conjugated anti-immunoglobulin at 1:1000 (IgG Alexa Fluor 594, 488, Invitrogen, Cat#:A-11001, A-11012). The microvasculature was stained using Lectin-594 conjugate (1:250, Vector Laboratories, Cat#:DL-1177-1). Images were obtained by taking a Z-stack containing 20μm thick sections. Images were taken at both 20×, 40×, and 63× magnifications using a Zeiss LSM900-Airy confocal microscope, with settings kept constant between respective image acquisitions used in quantification. For Iba1, NeuN, GFAP, and lectin staining, three coronal sections per mouse were analyzed.

### 2.12 Enzyme-linked immunosorbent assay (ELISA)

Plasma was separated from whole blood by centrifugation at 1500 x g for 15 min at 4 °C and together with supernatants from *in vitro* and brain and spleen homogenates from *in vivo* experiments stored at −80 °C until analysis. The protein concentration was determined using the bicinchoninic acid assay (BCA) (Fisher Scientific), and equal protein concentrations were standardized by diluting samples in PBS. Quantification of IL-6, TNF-α, and IL-10 was performed with the ELISA Mouse DuoSet kits (Bio-Techne Cat#:DY406, #DY410, and #DY417 ) with a sensitivity of 15.6 ρg/ml for IL-6 and 31.2 ρg/ml for IL-10, and TNF-α, respectively. Assays were performed following the manufacturer’s instructions, loading a total of 100µg of protein into each well in duplicate, and the optical density of each well was measured using a Multiscan FC microplate reader (Thermo Scientific) at 450nm.

### 2.13 Image processing and analysis

ImageJ (Fiji) was used to analyse the confocal images. To assess the mean intensity and area coverage by Iba1 and GFAP, 40× magnification stacks were merged, the background subtracted, and the image was converted to a binary image by threshold, and the area fraction was measured. GFAP images were taken at different anatomical locations in the brain, and to account for differences in cell density and morphology between regions, the data were normalized. Analysis of microglia morphology (process length and endpoints) was performed as previously described [41]. Briefly, an unsharp mask with a pixel radius of 3 and a mask weight of 0.6 value was used to increase contrast, followed by a despeckle step removing noise. The image was converted to a binary image by threshold and the steps despeckle, close- and removal of outliers (pixel radius 2, threshold 50) was applied to remove noise. The image was skeletonized using the skeleton plug-in and parameters extracted with “analyze skeleton 2D/3D”. Short branches were removed with a cut of value of 5. Settings were kept constant. For NeuN, the images were thresholded and cells counted by the “analyzing particles” plug-in with a cut size of 5μm. Settings were kept constant between analyses. The total number of cells was divided by the total field of view (F.O.V) area.

### 2.14 Statistical analysis

GraphPad Prism-9 was used for statistical analyses. Outliers were identified using the ROUT test, Q=1%. Normality was tested using the Shapiro-Wilk test. Ordinary one-way ANOVA or Kruskal-Wallis test was performed using Bonferroni or Dunn’s for multiple comparisons. Kaplan-Meier survival curves were analysed using a log-rank (Mantel-Cox) test. For comparison of *in vitro* ELISA data a two-way ANOVA and a Bonferroni correction for multiple comparisons was performed. Information about statistical tests and the number of animals is specified in the respective figure legends.

## 3. Results

### 3.1 Extracellular vesicles alleviate pneumococcal-induced neuronal cytotoxicity, and blockade of IL-6 trans-signalling mitigated inflammation

Firstly, we confirmed that the WT.EVs and the bioengineered EVs possessed the canonical tetraspanin EV surface markers CD9, CD63, and CD81 (**Figure 1A**), and the intravascular protein TSG101, but did not express calnexin, confirming purity (**Figure 1B**). In addition, the bioengineered EVs displayed RVG, IL-6, or both on their surface for RVG.EV, IL-6.EV and DB.EV respectively (**Figure 1A**). All EVs showed similar size profiles assessed through NTA and TEM (**Figure 1C-D**). The bioengineering of the EVs was therefore successful and did not alter the relative size or the expression of canonical proteins compared to WT.EVs.

**Figure 1.**
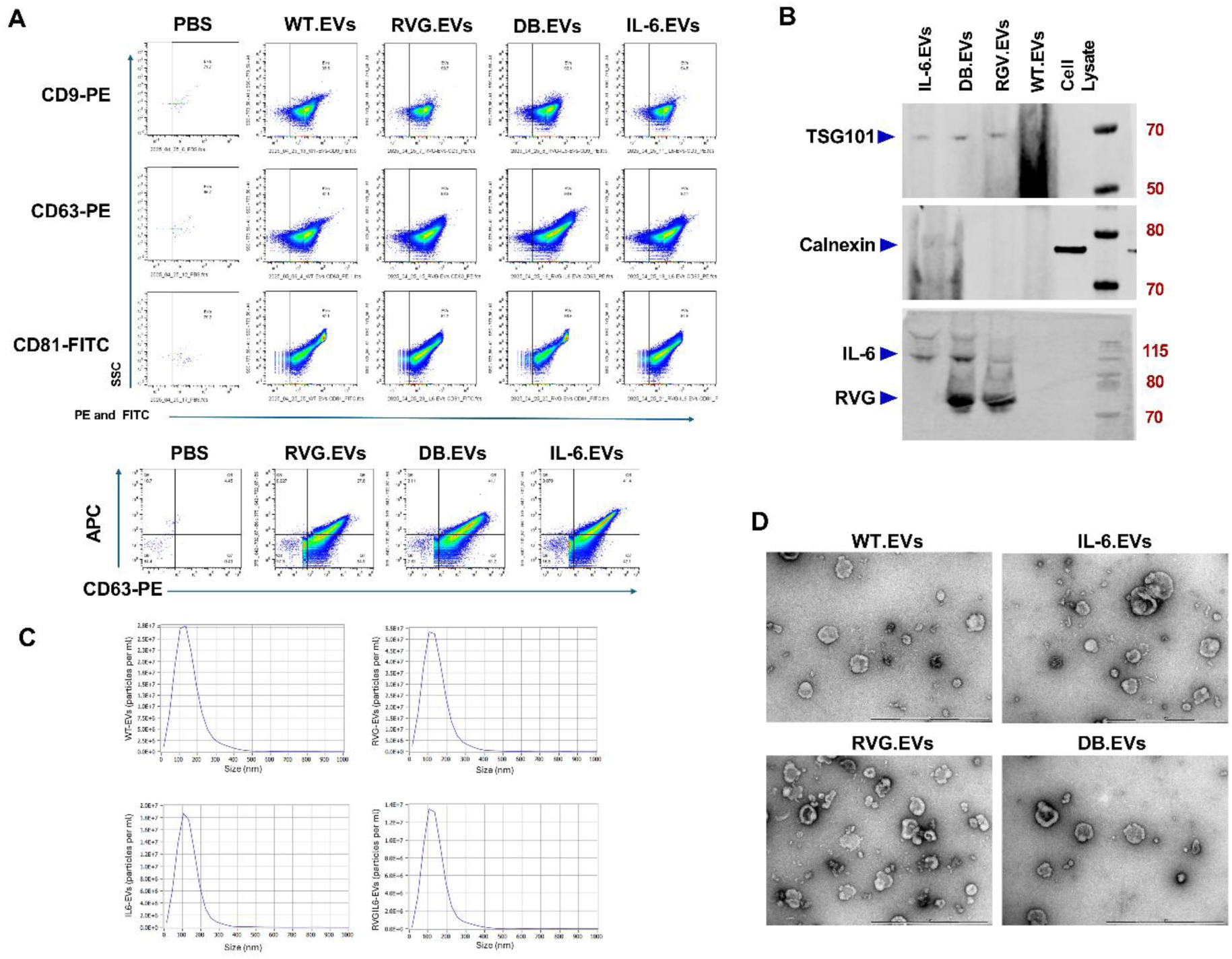
Characterization of WT.EVs and bioengineered EVs. The characterization of the EVs used in the *in vitro* and *in vivo* studies. (**A**) Flow cytometry analysis depicted in ***upper panel***: similar expression of the tetraspanin EV markers CD9 CD63 CD81, between the EV types. ***Lower panel***: confirmation of surface-exposed RVG peptide (APC-RVG) and IL-6ST (APC-IL-6ST), in RVG.EVs, DB.EVs and IL-6.EVs respectively. (**B**) Western blotting visualized the expression of IL-6ST and RVG peptide for the engineered EVs. The intravascular protein TSG101 was present in all EVs, while calnexin was used as a cellular organelle marker. (**C**) NTA was performed to assess the size of the EVs for the WT.EVs, RVG.EVs, IL-6.EVs, and RVG.IL-6.EVs (DB.EV). (**D**) Transmission electron microscopy for the WT.EVs and the engineered EVs, scale bar indicates 1µm.

To assess the potential of the EVs to block *S. pneumoniae* interaction with neurons, we infected differentiated human SH-SY5Y neuronal cells with T4 alone, in combination with WT.EV, RVG.EV, IL-6.EV and DB.EV, or with respective EV treatment applied 30 min post-infection. Presence of all EVs led to a significant reduction of bacterial adhesion (**Figure 2A**) and further reduced neuronal cytotoxicity (**Figure 2B**). Imaging revealed that the EVs reduced the adhesion of *S. pneumoniae* by covering the plasma membrane of the neurons (**Supplementary Figure 1A, B**). Importantly, treatment with EVs after infection also markedly reduced adhesion of the pneumococcus to neurons, and in combination with ceftriaxone treatment depicted a trend in reducing pneumococcal-induced cytotoxicity (**Supplementary Figure 2A, B**).

**Figure 2.**
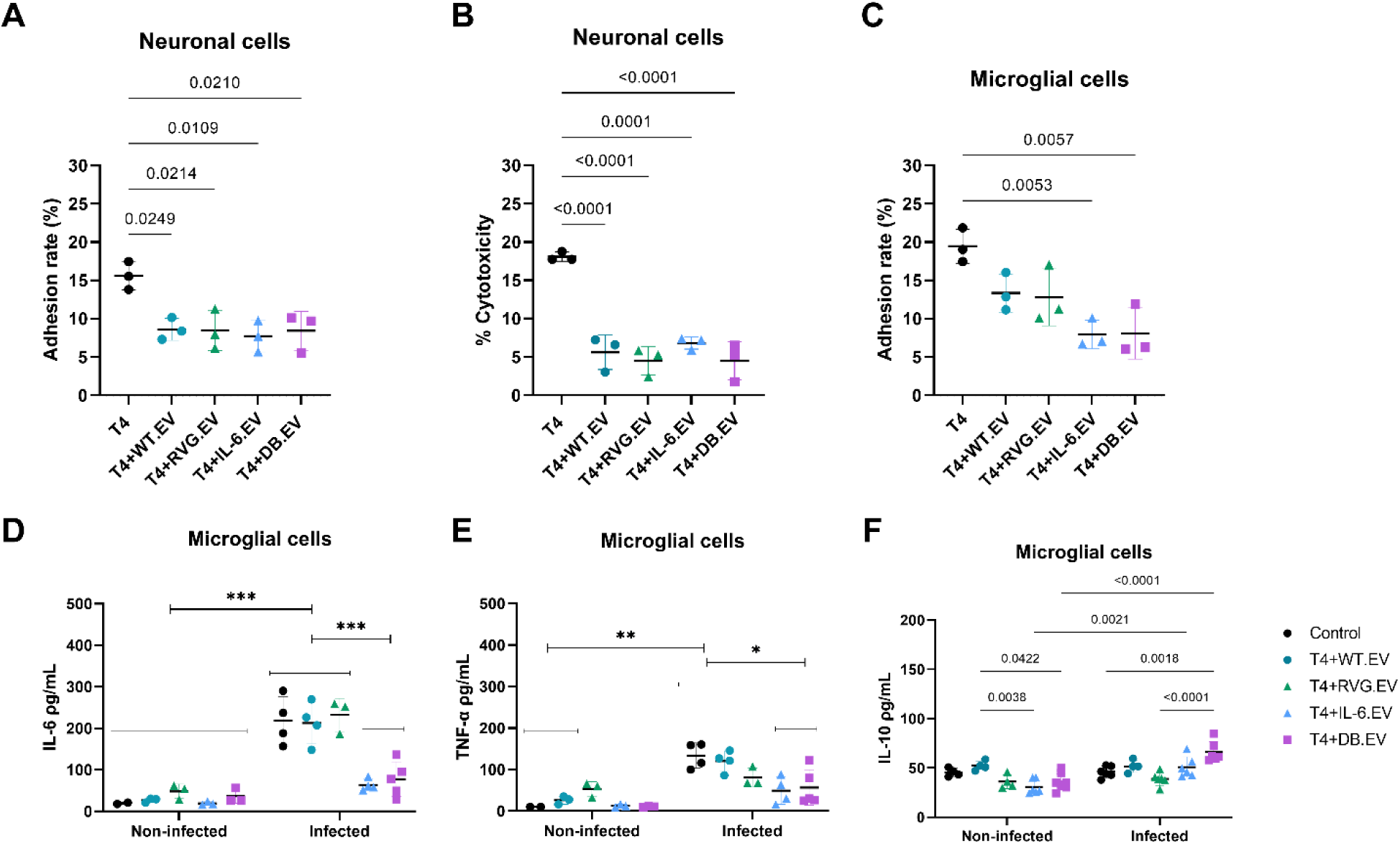
Treatment with EVs during *S. pneumoniae* infection had neuroprotective effects and reduced the release of pro-inflammatory cytokines. Differentiated SH-SY5Y neuronal-like cells infected with *S. pneumoniae* were treated with WT.EVs and bioengineered EVs, this treatment reduced *S. pneumoniae* adhesion rate (**A**) and cytotoxicity measured by LDH release (**B**). Treatment also reduced the adhesion of *S. pneumoniae* to BV-2 microglial cells (**C**). IL-6.EV and DB.EV treatment reduced the release of IL-6 (**D**) and TNF-α (**E**), while the release of IL-10 was increased after DB.EV treatment (**F**) during infection of microglial cells. For A-C, n=3 for all treatment groups, data were analyzed with a one-way ANOVA and Bonferroni post-hoc test for multiple comparisons. For D-F n=2-5. The data were analyzed using 2-way ANOVA with Bonferroni correction for multiple comparisons. Bars depict mean ± SD.

Next, we assessed how the EV treatment would impact inflammation in infected BV-2 microglial cells. We observed a reduction in bacterial adhesion upon treatment with EVs, with IL.6.EV and DB.EV treatments led to a significant reduction (**Figure 2C**), which was retained when EVs were added post-infection (**Supplementary Figure 2C**). Further, we assessed whether the level of cytokine release was altered during treatment with EVs in non-infected and infected microglia. The IL-6 levels and TNF-α levels were markedly reduced during infection for IL-6.EV and DB.EV-treated cells (**Figure 2D, E**). BV-2 microglial cells upregulated the anti-inflammatory cytokine IL-10 in response to LPS and CNS injury [42, 43]. Thus, we measured IL-10 release, which upon infection, was increased for the DB.EV (**Figure 2F**). When applying the EV treatment 30 min post-infection, no differences was observed in cytokine level, however in combination with ceftriaxone treatment IL-6.EV treatment non-significantly reduced TNF-α levels (**Supplementary Figure 2D**). These results demonstrate the neuroprotective effect of the EVs, and that expression of the signal incompetent IL-6ST decoy receptor reduced inflammation during infection in neurons and microglia *in vitro*.

### 3.2 EVs sequester Ply, reducing its cytotoxicity

We hypothesized that the EVs, rich in cholesterol, could sequester Ply and thereby neutralize the toxin. To test this, 2.2 x 10^11^ p/ml EVs were incubated with 2.0µg/ml of recombinant Ply. All EV types bound Ply (**Figure 3A, B**), with no difference observed between the size of EVs during incubation (**Supplementary Figure 3**). Although not statistically significant, DB.EV depicted a trend towards a higher concentration of Ply in the supernatant, indicating reduced sequestration capacity (**Supplementary Figure 4A,B**). In order to confirm that this sequestering caused a neutralization of the toxin, the hemolytic activity of Ply, which induce lysis of red blood cells at various concentrations (**Supplementary Figure 4C**) were assessed upon treatment with different concentrations of WT.EV. WT.EVs reduced 1.0µg/ml Ply induced lysis at 1 x 10^8^ p/ml (**Supplementary Figure 4D**). However, the strongest effect was observed at a concentration 1 x 10^9^ p/ml, and at this concentration all EV groups significantly reduced the hemolytic activity of Ply, confirming their neutralization capacity (**Figure 3C).** To confirm that this neutralization also protected against Ply-induced cytotoxicity towards cells, we treated SH-SY5Y neuronal cells with either 1μg/ml (**Figure 3D**) or 2μg/ml (**Figure 3E**) Ply in combination with all the EV types and assessed LDH release as a measure of cytotoxicity. All EV treatments reduced the cytotoxicity of Ply. Taken together, these results demonstrate that EV treatment can protect against the cellular toxicity induced by Ply. Next, we sought to evaluate the therapeutic potential of the EVs *in vivo*.

**Figure 3.**
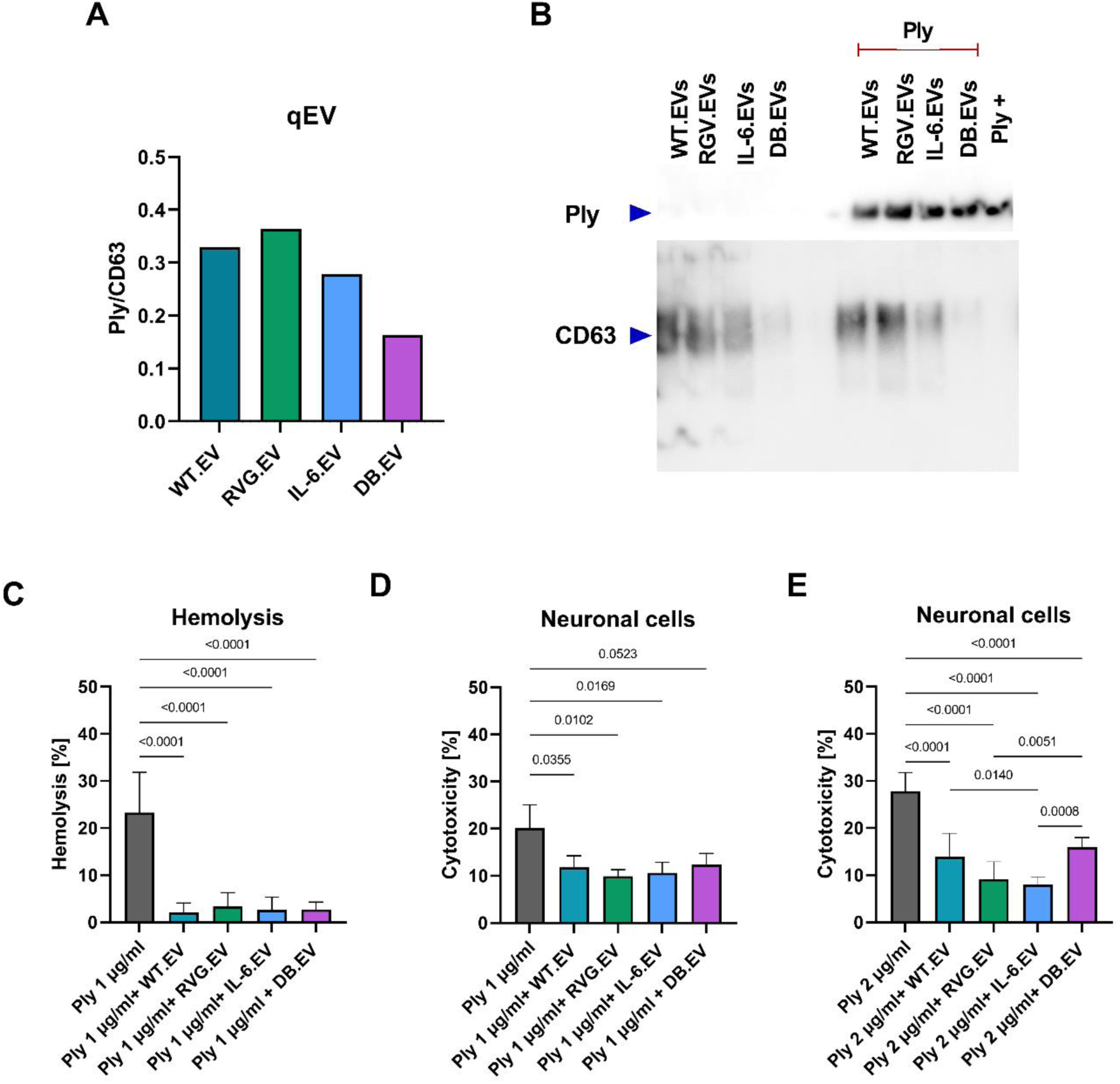
WT.EVs and bioengineered EVs sequestered and neutralized Ply. Ply sequestration was confirmed by incubating 2μg/ml of Ply with 1×10^11^ p/ml EV followed by SEC with qEV columns and WB analysis of elute. Quantification demonstrated that all EVs sequestered Ply, graph shows the ratio of band intensity measured by AUC of Ply/CD63 (**A)** (**B**) Western blot for Ply and CD63 were used for the quantification Fig3A. (**C**) The hemolysis of red blood cells was analyzed upon treatment with Ply 1μg/ml, and the neutralization capacity of the EVs was demonstrated for all EV groups. Differentiated SH-SY5Y neuronal cells were exposed to 1 or 2μg/ml of Ply and treated with 1×10^9^ p/ml of respective EVs, which significantly reduced cytotoxicity (**D**, **E**). qEV experiment was performed once, while n=3 for C, D, and 4 for E. The data was analyzed by a one-way ANOVA, with Bonferroni correction for multiple comparisons. Bars depict mean ± SD.

### 3.3 Treatment with extracellular vesicles increased survival during experimental pneumococcal meningitis

First, to determine if the EVs confer a reduction in *S. pneumoniae* growth *in vivo*, and to confirm that the EVs could enter the brain, we tested three different doses of WT.EVs: 0.5,1, and 2 x 10^11^ p/mouse in our established bacteremia-derived pneumococcal meningitis model [39]. Mice were infected with

*S. pneumoniae* intravenously and received EV treatment one hour later, control animals received PBS. At 8 hours post-infection, no differences in bacterial load in the brain, spleen, or blood were observed between controls and WT.EV treatment groups (**Supplementary Figure 5A**). Importantly, we confirmed that the WT.EVs successfully entered the brain parenchyma (**Supplementary Figure 5B**). Given the *in vitro* potency of DB.EV treatment, we preliminarily assessed whether a dosage of 2 x 10^11^p/mouse of DB.EVs would have an effect *in vivo.* The mice were sacrificed upon developing severe symptoms of pneumococcal infection. We confirmed that DB.EV treatment significantly improved clinical outcome and increased survival (**Supplementary Figure 5C, D**), validating its therapeutic potential and justifying a full-scale study.

Subsequently, all EV treatment groups were evaluated (**Figure 4A**). In addition, to investigate the early brain pathology and the dynamic inflammatory response observed during pneumococcal meningitis [44], five DB.EV-treated mice were sacrificed at the same time point as the PBS-treated infected control group (T4), to minimize time as a confounder in our downstream analysis. The infected control group (T4) had a median time of survival of 13 hours; at this time, the DB.EV group presented mild symptoms and was termed “DB.EV mild” (**Figure 4B**). All other groups were sacrificed when developing severe symptoms of pneumococcal disease.

**Figure 4.**
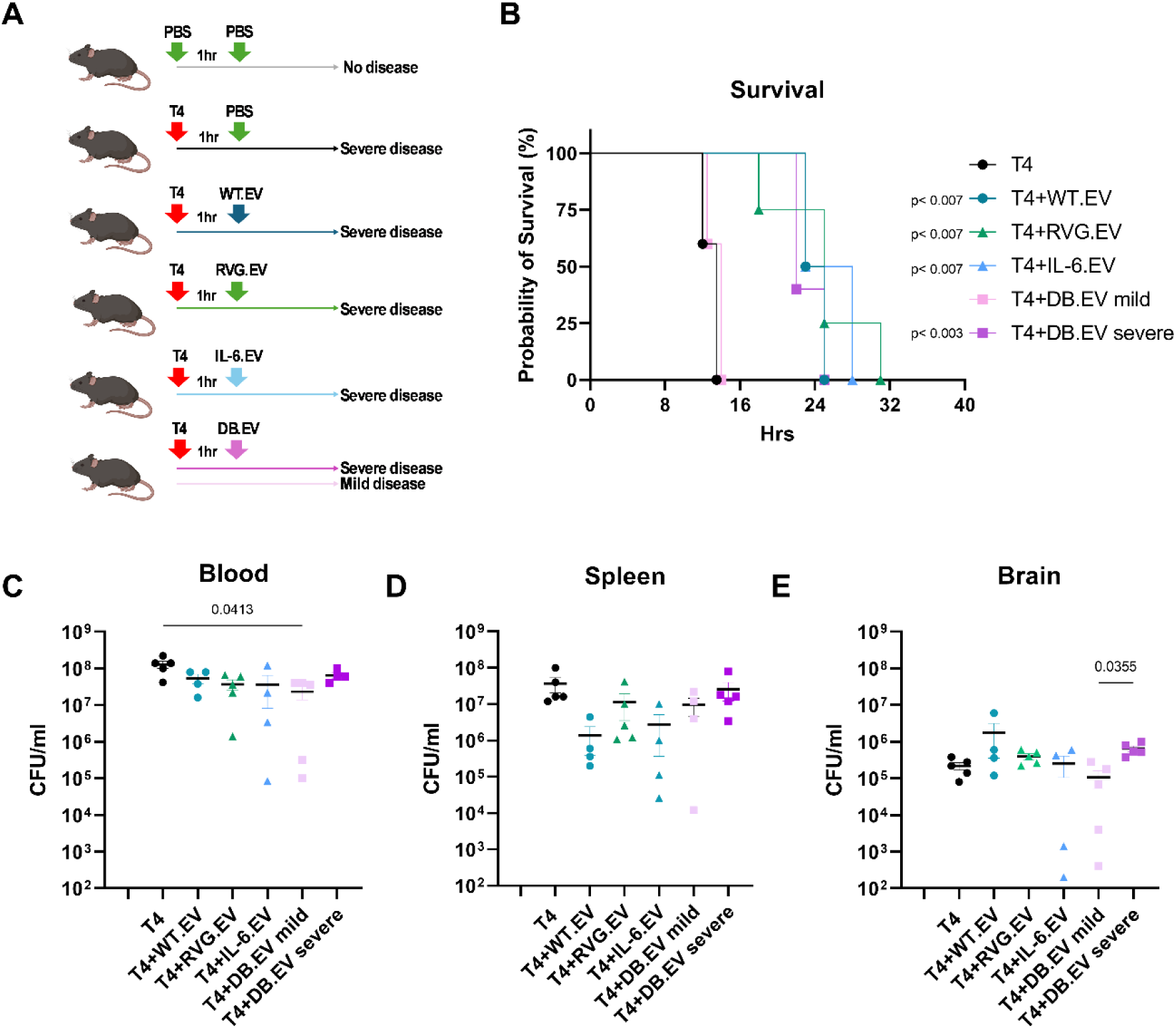
Treatment with EVs increased survival without affecting bacterial loads. Experimental overview of animal groups and treatments (**A**). Treatment with EVs significantly increased the survival (**B**). At the time of the experimental endpoint, the animals that reached severe infection showed no significant differences in their bacterial load in the blood (**C**), spleen (**D**) or brain (**E**). DB.EV mild animals showed a slight reduction in bacterial load in the blood and brain. n= 4 for WT.EV and IL-6.EV treated groups, n= 5 for T4, RVG.EV, DB.EV for all graphs. Data were analyzed using a one-way ANOVA and Bonferroni correction for multiple comparisons. Survival curve comparison was tested with the Mantel-Cox test, the respective p-value in comparison with T4 (control infected) is annotated next to each group’s legend. Bars depict mean ± SD.

All EV treatments significantly improved symptomatology and survival, with a median time of survival of 24 hours compared to infected controls (T4) (**Figure 4B**). The bacterial load in the blood (**Figure 4C**), spleen (**Figure 4D**), and brain (**Figure 4E**) did not differ between severe symptom groups; however, the DB.EV mild group had slightly lower bacterial numbers in the blood compared to infected controls (T4), and in the brain compared to the DB.EV severe animal group (**Figure 4C**). Our results confirmed the therapeutic efficacy of EVs in experimental pneumococcal meningitis, with Ply sequestering as the putative mechanism for the delayed disease onset, due to different EV treatments evoking similar beneficial therapeutic effects. We next assessed if the bioengineered EVs affected the host inflammatory response.

### 3.4 Treatment with RVG.EVs reduced inflammation in the periphery and brain

The inflammatory response is the main driver of cellular damage in pneumococcal infections, especially in meningitis. Given that the expression of the signal incompetent IL-6ST decoy receptors reduced the pro-inflammatory response during infection *in vitro*, we assessed whether treatment with EVs modulated the cytokine release *in vivo*.

In the spleen, a crucial component of the immune system’s periphery, we observed an increased release of IL-6 upon infection in the T4 group compared to non-infected controls (**Figure 5A**). This increase was significantly reduced for all EV-treatment groups, with the RVG.EV and IL-6.EV groups restoring IL-6 levels close to baseline. TNF-α release was also elevated upon infection and the WT.EV-treated animals depicted the highest levels. Importantly, all bioengineered EV treatments reduced the levels of TNF-α (**Figure 5B**). In the brain, the DB.EV severe group had the highest expression of both IL-6 and TNF-α, however, also the WT.EV animals had increased levels of both cytokines (**Figure 5C, D**). In contrast, the RVG.EV-treated group had significantly lower expression. Additionally, in a separate *in vivo* study, we evaluated the effect of RVG.EV and IL-6.EV treatment on blood cytokine levels, both treatments reduced IL-6 and TNF-α compared to T4 controls (**Supplementary Figure 6A-E**). These findings suggest that individual EV treatments modulated inflammation differently, with the RVG.EVs depicted the best neuroprotective effect.

**Figure 5.**
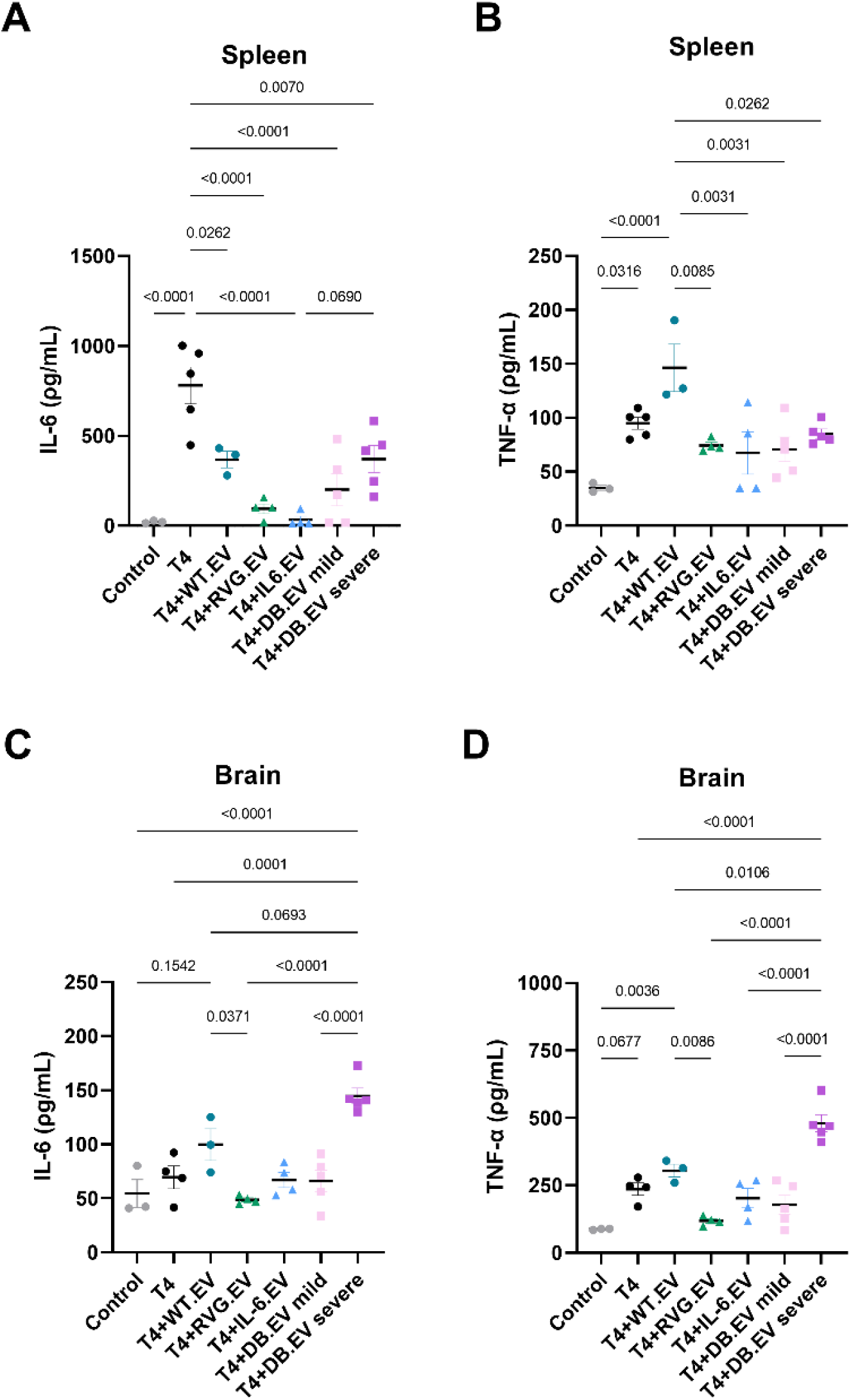
EV treatment altered the release of pro-inflammatory cytokines in the spleen and brain during infection. In the spleen, the T4 group had the highest levels of IL-6, however treatment with EVs significantly reduced the release, with the IL-6.EV group depicting the lowest level (**A**). TNF-α was also significantly increased during infection, however treatment with the bioengineered EVs significantly lowered the levels of TNF-α (**B**). In brain homogenates, the levels of IL-6 (**C**), and TNF-α (**D**) were increased during infection for the animal groups with only the RVG.EV depicted lower levels. n= 3 for control and WT.EV, n=4 for RVG.EV and IL-6.EV, n=5 for DB.EV animal groups. Data were analyzed using a one-way ANOVA and Bonferroni correction for multiple comparisons. Bars depict mean ± SD.

### 3.5 IL-6ST-expressing EVs modulate glial activation without affecting neuronal survival

To assess if treatment of pneumococcal meningitis with different EVs impacted the cellular response in the brain, we analyzed changes in the microglia, astrocytes, and neuronal population. All infected groups had microglia depicting morphological changes compared to the non-infected controls (**Figure 6A and Supplementary Figure 7A *upper panel*).** Infection decreased both microglial process length (**Figure 6B**) and endpoints (**Figure 6C**). However, the RVG.EV and DB.EV mild treated groups, demonstrated a significant increase in process length compared to the T4 control group (**Figure 6B**), indicating reduced microglial activation. Furthermore, both groups, in addition to the IL-6.EV-treated group had significantly increased endpoints compared to the other infected groups (**Figure 6C**). Interestingly, a significant increase in Iba1 intensity was only observed in the IL-6.EV and DB.EV severe-treated group (**Figure 6D**). In addition, both groups showed a lower area coverage of GFAP signalling, indicating reduced astrocytic coverage and protrusion retraction (**Supplementary Figure 7A *middle panel*, Figure 6E**). To assess if there was a difference in the number of neurons in the cortex, we stained with the neuronal cell body marker NeuN (**Supplementary Figure 7A *lower panel***), we observed no statistical differences between groups, however, the WT.EV-treated animals had a slight trend of reduced numbers of neurons and intensity of staining (**Supplementary Figure 7B, C**). In summary, RVG.EV treatment reduced the microglial morphological changes, while the IL-6 trans-signalling likely plays a key role in brain cellular communication during infection, and its blocking may disrupt glial function, without though exacerbating neuronal loss during the acute phase of infection.

**Figure 6.**
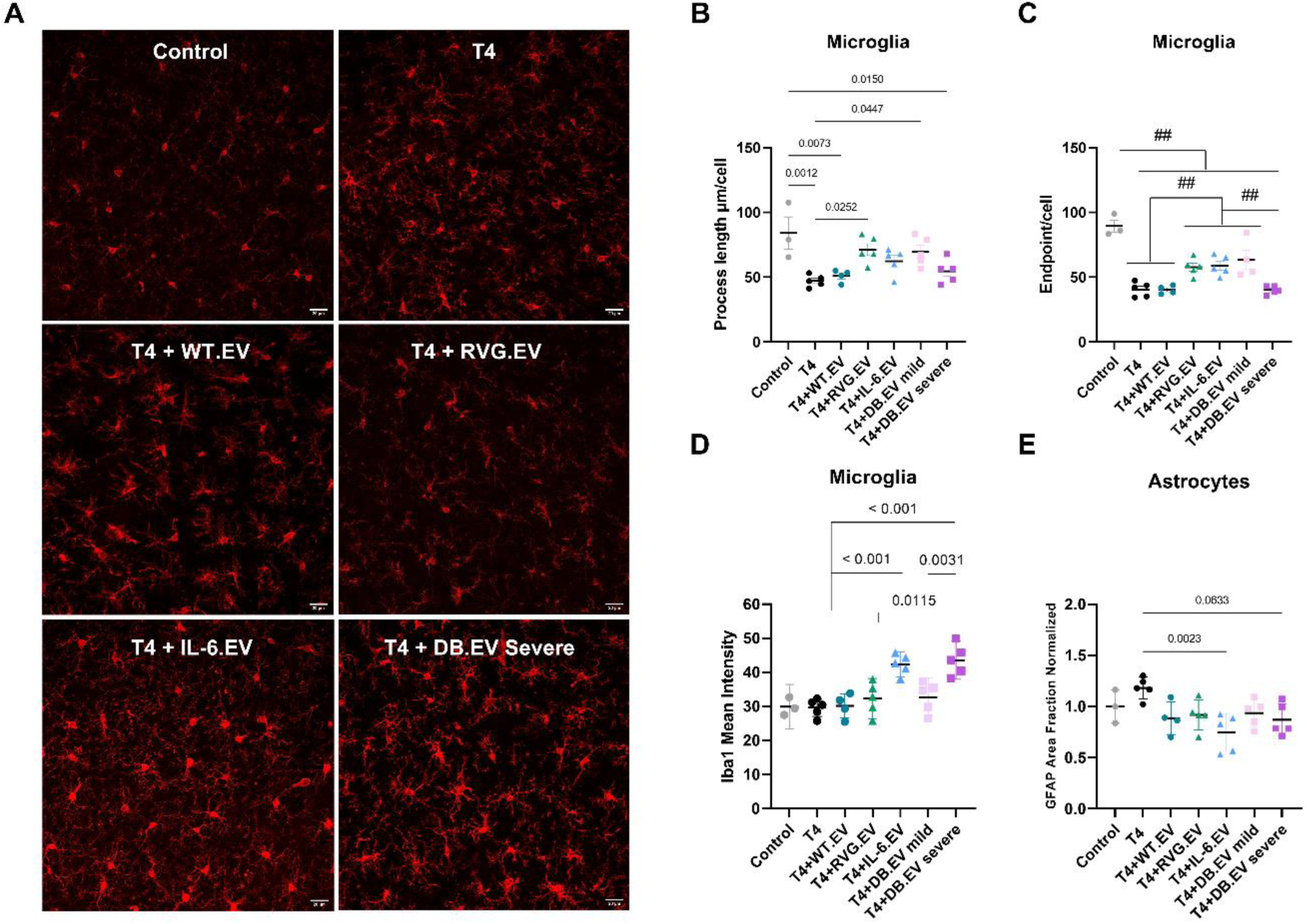
IL-6ST decoy receptor expression increased microglial activation and reduced GFAP signal, while RVG.EV treatment indicate a protective effect. Representative images from immunofluorescence analysis (**A**) of microglial cell population (Iba1, red), scalebar depicts 20μm In-depth analysis of microglial morphology demonstrated that T4 infection reduced microglial process length (**B**) and endpoints (**C**), with different EV treatment affecting both processes. The mean intensity signal of Iba1 (**D**) and GFAP area coverage (**E**) were significantly increased for IL-6.EV and DB.EV animal groups compared with controls and the other infection groups. n= 3 for control, n=4 for WT.EV, and n=5 for RVG.EV, IL-6.EV and DB.EV animal groups. Data was analyzed using a one-way ANOVA and Bonferroni correction for multiple comparisons. Bars depict mean ± SD

## 4. Discussion

Bacterial meningitis, frequently caused by the pneumococcus, is a devastating neurological disease, which despite the availability of antimicrobials and pneumococcal vaccines, the incidence remains high [45]. Survivors have a high risk of developing long-term neurological sequelae, which are consequences of neuronal damage caused by neuroinflammation and released bacterial compounds, such as Ply [10, 15]. The treatment strategy is complex, as patients often suffer from complications, such as sepsis and myocarditis, but also brain abscesses and cerebrovascular disease [46–48]. The low capacity of antibiotics to cross the BBB and the often cytolytic nature of these render the brain sensitive to the bacteria and bacterial components, such as Ply, for a prolonged period [49]. It is therefore crucial to develop new therapeutic strategies that can efficiently reach the brain, reduce neuroinflammation, and protect resident brain cells. In this sense, the sequestering and neutralization of Ply is essential. In this study, we have, for the first time, evaluated the treatment potential of bioengineered EVs in pneumococcal meningitis. We have demonstrated that treatment with EVs increased the survival of the infected mice, primarily by sequestering Ply. In addition, by bioengineering the EVs to express RVG peptides, the protective effect of the EVs was extended to the brain, where the treatment reduced inflammation and demonstrated neuroprotective capacity.

EVs have a high therapeutic potential in pneumococcal meningitis and have already been successfully applied in both patients and experimental animal models for cancer, neurodegenerative disease, and systemic inflammatory disease [50, 51]. Depending on their source, EVs exhibit differences in their intrinsic properties, including low immunogenicity and high tissue penetration [31]. Further, the ability to engineer EVs to express and carry specific cargo increases their potential as therapeutic agents in a variety of diseases [52]. EVs are fast-acting molecules, and their translocation to tissue has been observed as early as a few minutes after administration [53]. In addition, they have a wide array of biodistribution and can exert their effects in multiple tissues. This wide biodistribution is particularly valuable in pneumococcal meningitis, where bacteria frequently invade peripheral organs [2, 34]. The crossing of neural barriers by native EVs involves transcytosis, and while no direct evidence is observed in humans, several *in vitro* models have demonstrated that EVs cross endothelial cell barriers in an inflammatory milieu [29, 30]. While using a human endothelial-cell monolayer can provide useful preliminary insights into drug delivery across the BBB, *in vitro* cellular systems remain an artificial model that lacks the full complexity and physiological context of the human BBB [54]. Thus, *in vivo* BBB passage data are essential to validate delivery strategies, meaning findings in animal models, such as the bacteremia-derived meningitis model used in this study. In pneumococcal meningitis patients, bacterial components and the systemic immune response cause significant vascular changes in the brain, including BBB disruption, facilitating an increase in antibiotic infiltration into the brain [5, 55]. However, treatment with steroids, such as dexamethasone in the case of pneumococcal meningitis, can reduce CSF antibiotic concentrations by limiting BBB permeability [56]. In this sense, EVs bioengineered with RVG peptides have a therapeutic advantage as they can cross human endothelial cell layers through receptor-mediated endocytosis, due to the presence of AchR on the BBB [34, 57]. The fact that the human BBB is more leaky than the mouse BBB in brain disease [58, 59] further underscores the strong translational potential of the engineered EVs used in our study. Their demonstrated ability to cross the BBB and infiltrate the brain during *S. pneumoniae* infection *in vivo* suggests that these EVs could be particularly promising for clinical applications in human patients, where BBB permeability during disease may further facilitate targeted therapeutic delivery.

During pneumococcal meningitis, host cells are exposed to the cholesterol-binding cytolysin Ply, which causes a plethora of harmful effects on the host, including cell lysis, activation of pro-inflammatory cascades, and increased production of reactive oxidative species [60, 61]. The neutralization of Ply during infection is crucial for reducing neuronal damage and, consequently, the onset of neurological sequelae in patients [60, 62]. Previous studies using liposomes have demonstrated the interaction between cholesterol and Ply, with an increase in cholesterol content associated with increased sequestration. Similar to liposomes, the EV membrane contains cholesterol [28], and in our study, the EVs sequestered and neutralized Ply, which protected against Ply-induced neurotoxicity. Importantly, during this sequestration, the abundance of circulating Ply targeting host cells is reduced. *In vivo*, this ultimately mitigated both cytotoxicity and inflammation, delaying disease onset. The observed extension of survival was primarily due to the sequestration of Ply in the periphery, since all EVs delayed disease onset, with the RVG.EV treatment most effective in the brain.

Ply induces a pro-inflammatory response and contributes to the peripheral and neuroinflammatory process during pneumococcal meningitis; the latter is the main cause of neurological sequelae in patients [13, 63]. Microglia, together with other brain resident cells, recognize bacterial compounds inducing upregulation of the NF-κβ pathway and the release of pro-inflammatory cytokines [10]. Dexamethasone treatment reduces inflammation in patients and has been reported to improve outcomes; however, animal studies indicate that an inflammatory response promotes survival [64, 65]. It exists a very delicate balance between beneficial and harmful inflammatory responses during pneumococcal meningitis; an inadequate response induces a higher risk of tissue damage [66–68], while prolonged inflammation increases tissue damage and risk of death [10]. Increased levels of TNF-α and IL-6 correlate with tissue damage in animal models, and these cytokines are markedly increased in the CSF of bacterial meningitis patients [69, 70]. IL-6 is produced in response to infection and cellular damage and exerts its pro-inflammatory effects by binding IL-6ST receptors ubiquitously expressed, termed the trans-signalling pathway [71]. Treatment with EVs expressing signal incompetent IL-6ST decoy receptor reduced LPS-induced systemic inflammation and neurodegenerative disease pathogenesis [35, 72]. In our study, EVs expressing signal-incompetent IL-6ST decoy receptors (IL-6.EVs) reduced peripheral IL-6 levels yet failed to dampen inflammation in the brain. Moreover, they induced increased microglial activation and astrocytic process retraction. We can only hypothesize that the trans-signalling pathway of IL-6 plays an important role in cellular communication in the brain during infection, and its absence alters the function of glial populations. However, the effect on the glial populations did not cause neuronal damage.

Interestingly, treatment with RVG.EV significantly lowered TNF-α and IL-6 levels in the brain and the periphery. While the protective effect of the RVG.EV treatment in the brain could be explained by its higher affinity for brain infiltration, where they neutralize and sequester Ply: the anti-inflammatory effect in the spleen and blood was not expected. The spleen has the second-highest uptake of EVs in the body after the liver, and it might cause an increased Ply sequestration and reduced inflammation [73]. Additionally, the spleen, while not directly innervated by the vagus nerve, plays a crucial role in the systemic cholinergic anti-inflammatory pathway through the splenic nerve and abundance of Ach receptors [74]. The binding of Ach to these receptors has an inhibitory effect on inflammation [75]. Although the RVG peptide has been demonstrated to be a competitive antagonist using the *Xenopus laevis* oocyte model, it is to our knowledge no evidence on the effect of the RVG peptide on immune cells [76]. The overall anti-inflammatory effect of RVG.EV treatment in the *in vivo* study was not observed for the microglial cell line, highlighting the complexity of the immune response during pneumococcal meningitis.

In this study, we have exclusively focused on EV treatment in pneumococcal meningitis, which is the most severe form of pneumococcal disease. The neutralization of Ply by the EVs will also apply to other pneumococcal infections, such as myocarditis and pneumonia, where Ply contributes to the damage on cardiac and lung cells, respectively [77, 78]. Our results support previous research using liposomes, that neutralization of Ply is key to the successful treatment of pneumococcal infections [79–81]. Additionally, cholesterol-binding cytolysin production is not a unique feature of the pneumococcus; other meningitis-causing bacteria also produce this type of toxin, including *Streptococcus pyogenes,* which produces Streptolysin O [82]. Ultimately, using EVs to reduce toxicity can also be extended to these infections. The therapeutic potential of EVs, liposomes, and other Ply neutralizing agents is especially relevant in combination with bacteriolytic antibiotics. These antibiotics cause bacterial lysis, augmenting Ply release and increasing tissue damage [83]. Also relevant for the potential therapeutic feasibility of the RVG.EVs are the potential adverse effect of the RVG peptides on cholinergic neurons. The peptide is crucial for uptake of the rabies virus into neurons and can influence neuronal excitability and signalling; causing behavioural changes in model systems [76, 84]. However, RVG exosome treatment remains a promising strategy for targeting CNS disease, with previous studies demonstrating no adverse effect of the treatment [85]. Multiple preclinical studies engineered exosomes with RVG (Lamp2b–RVG or RVG29 surface display) to deliver siRNA or therapeutic cargos to the brain after systemic delivery; these studies reported efficient neuron targeting and gene knockdown and observed no non-specific uptake or attenuation on repeat dosing (no overt adverse effects), and subsequent preclinical work has reproduced effective brain delivery and therapeutic benefit without obvious acute toxicity [85, 86].

One important aspect to consider is that we evaluated EVs only during the acute phase of meningitis and as a standalone treatment *in vivo*. The bacteremia-derived mouse model, while reflective of natural infection routes, also induces sepsis, unlike models using direct CSF inoculation (Intracisternal (I.C) model) combined with antibiotics [87]. Nevertheless, the peripheral immune system and tissues are also involved in disease progression in most cases of pneumococcal meningitis [2]. In contrast to I.C models, the animals are sacrificed at an earlier stage in the bacteremia-derived model; thus, there are fewer neuropathological features [88]. While all EVs protected against pneumococcal interaction and Ply cytotoxicity *in vitro*, our *in vivo* results did not demonstrate a significant neuroprotective effect, potentially due to the high systemic disease burden in our animal model. Although we clearly showed that, *in vitro,* EVs can protect neuronal cells from bacterial interaction and Ply-induced cytotoxicity.

In conclusion, our results strongly underline the importance of Ply neutralization in pneumococcal meningitis and the effectiveness of EVs in this manner. Our results indicate that expression of RVG peptides on EVs had a substantial benefit as it suppressed peripheral inflammation, increased the EVs capacity to reach the brain, where they sequestered Ply; consequently, lowering neuroinflammation and protecting against *S. pneumoniae*-induced cytotoxicity. Taken together, RVG-expressing EVs present a compelling adjuvant strategy for future treatment of pneumococcal meningitis.

## Data availability

All data generated and analyzed in this study are included or can be provided upon request.

## Author contributions

KF, MVT, and FI designed the study, DRM, XL, SEA, and OPBW contributed to the experimental approach, DRM, XL, HZ VWQH, SEA, and OPBW engineered and performed the characterization of the EVs. KF and MTV performed data collection and analysis, all authors contributed to the interpretation of the results. KF drafted the manuscript. All authors critically revised the manuscript.

