## Supplementary Material for "Bioengineered extracellular vesicles mitigate neuroinflammation by neutralizing pneumolysin and delaying disease onset in experimental pneumococcal meningitis"

| **Study** | **1** | **2** | **3** | **4** |
|---|---|---|---|---|
| **Type** | Pilot | Pilot | Full-scale | Add-on |
| **Purpose** | • Confirm dosage safety and titration<br>• *S. pneumoniae* growth in presence of EVs<br>• EV crossing of the BBB | • Assess DB.EV effect on survival | Assess EV treatment effect on:<br>• Survival<br>• CFU<br>• Inflammation<br>• Neuroprotection | Assess EV treatment effect on:<br>• Blood cytokine levels |
| **Mice (n)** | 3 each group | 4-7 | 3-5 | 3 |
| **EV type** | WT.EV | DB.EV | WT.EV, RVG.EV, IL-6.EV, DB.EV | RVG.EV, IL-6.EV |
| **Endpoint and tissue collection** | 8 hours post-infection | Score of 0.4/0.5 | Score of 0.4/0.5*<br>*DB.EV mild sacrificed together with T4 control | Score of 0.4/0.5 |
| **Downstream analysis** | CFU, WB for CD81 | Survival, Symptoms | CFU, cytokine release, immunohistochemistry | Cytokine release |
| **Figure** | Supplementary Figure 4A.B | Supplementary Figure 4C.D | Figure 4-6, and Supplementary Figure 7 | Supplementary Figure 6 |

**Supplementary Table 1.** Overview of the four *in vivo* studies included, their purpose, n-number, type of EV used, endpoint and tissue collection, downstream analysis and the figure presenting the results.

| Comparison | p-value |
|---|---|
| Control vs. T4 | <0.0001 |
| Control vs. T4+WT.EV | <0.0001 |
| Control vs. T4+RVG.EV | 0.0002 |
| Control vs. T4+IL-6.EV | 0.0002 |
| Control vs. T4+DB.EV mild | 0.0033 |
| Control vs. T4+DB.EV severe | <0.0001 |
| T4 vs. T4+RVG.EV | 0.0315 |
| T4 vs. T4+IL-6.EV | 0.0181 |
| T4 vs. T4+DB.EV mild | 0.0033 |
| T4+WT.EV vs. T4+RVG.EV | 0.0456 |
| T4+WT.EV vs. T4+IL-6.EV | 0.0272 |
| T4+WT.EV vs. T4+DB.EV mild | 0.0052 |
| T4+RVG.EV vs. T4+DB.EV severe | 0.0248 |
| T4+IL-6.EV vs. T4+DB.EV severe | 0.0142 |
| T4+DB.EV mild vs. T4+DB.EV severe | 0.0026 |

**Supplementary Table 2.** P-values for Figure 6C (microglial endpoints).

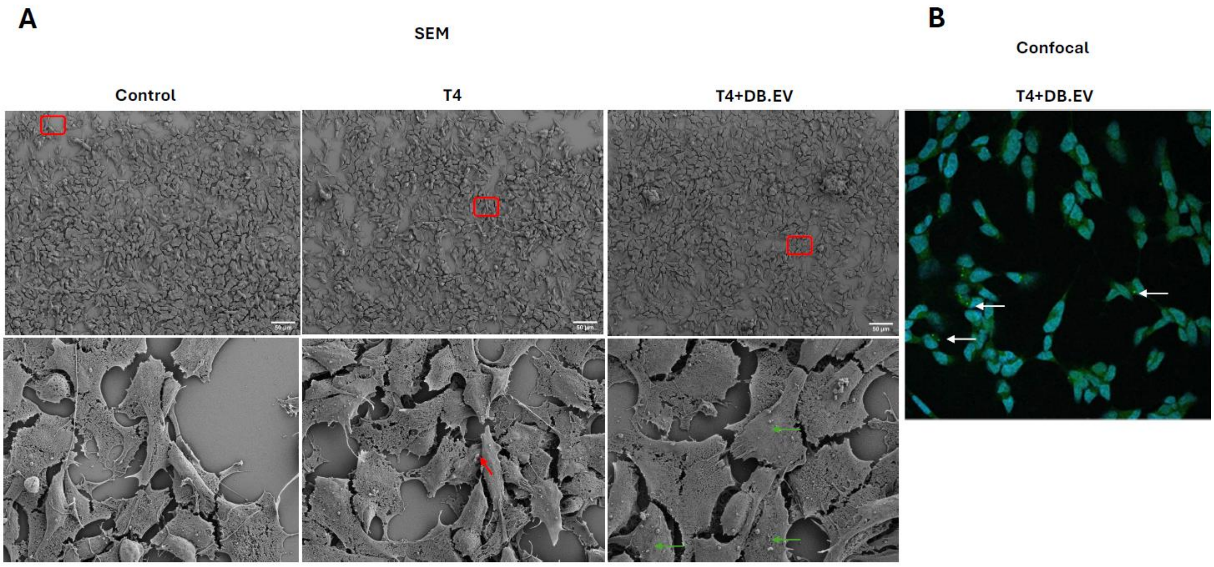


**Supplementary Figure 1.** Representative images from SEM showing control infected (T4), and infected + DB.EV treatment of SH-SY5Y neuronal-like cells (**A**). ***Upper panel*** indicates low magnification; red squares indicate the area of higher magnification in the ***lower panel***. Red arrows indicate *S. pneumoniae*. Green arrows indicate DB.EV clusters. Scalebar indicate 50µm The DB.EV was also observed targeting the SH-SY5Y neuronal cell body. representative image depicts confocal imaging (**B**). The white arrows indicate the EVs. represented as bright green circles.

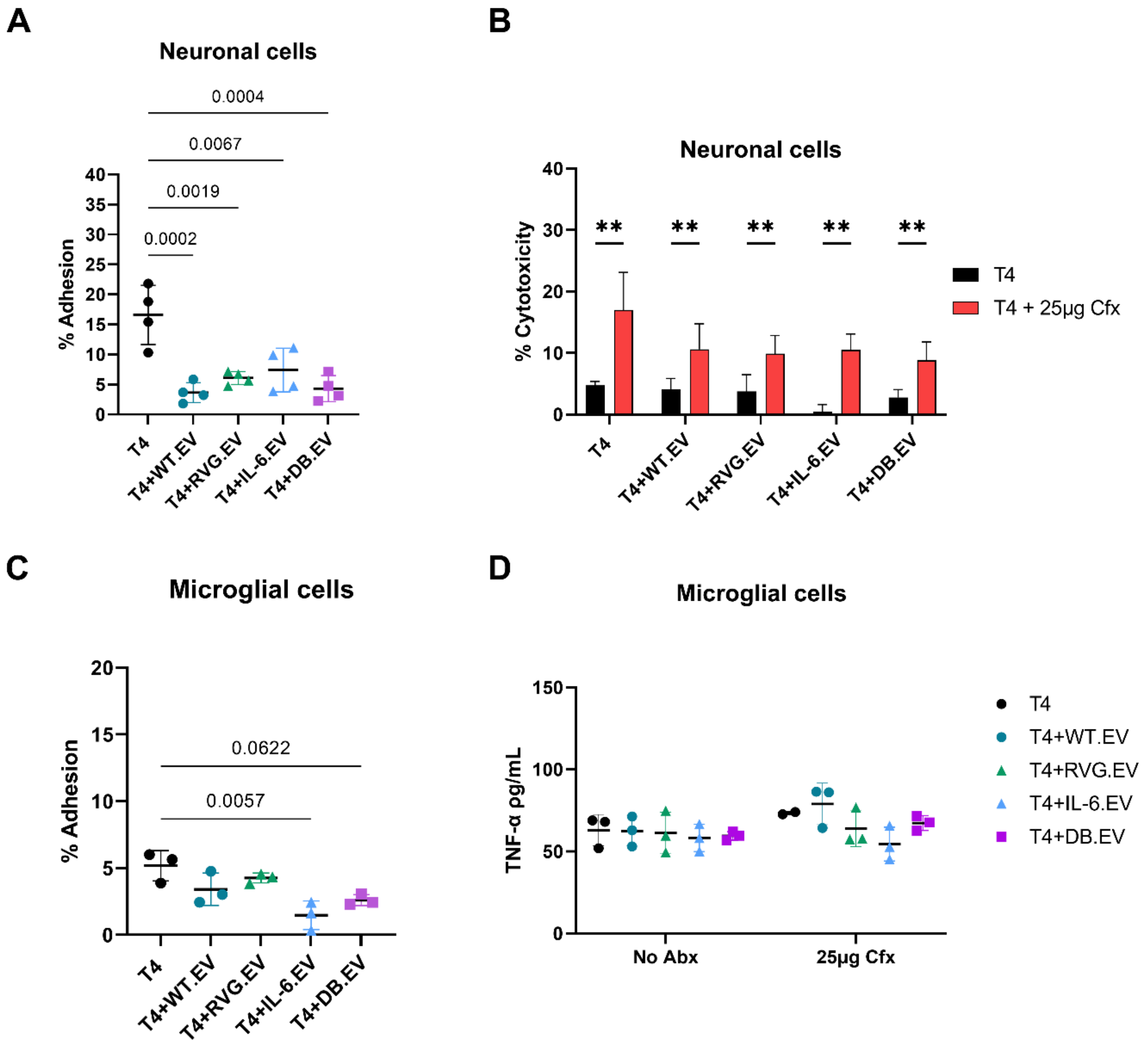


**Supplementary Figure 2. Treatment with EVs post *S. pneumoniae* infection with and without antibiotics demonstrated neuroprotective effects.** Differentiated SH-SY5Y neuronal-like cells infected with *S. pneumoniae* were treated with WT.EVs and bioengineered EVs 30 min post-infection, this treatment reduced *S. pneumoniae* adhesion rate (**A**). In combination with 25µg/ml Ceftriaxone (Cfx), the EV treatment demonstrated trend in reducing cytotoxicity measured by LDH release (**B**). Treatment also reduced the adhesion of *S. pneumoniae* to BV-2 microglial cells when added 30-min post infection (**C**). In combination with 25µg/ml Cfx, the EV treatment demonstrated trend in reducing IL-6.EV treatment demonstrated a trend in reducing TNF-α levels (**D**). For A-D, n=2-4 for all treatment groups, data were analyzed with a one-way ANOVA and Bonferroni post-hoc test for multiple comparisons. The data were analyzed using 2-way ANOVA with Bonferroni correction for multiple comparisons. Bars depict mean ± SD.

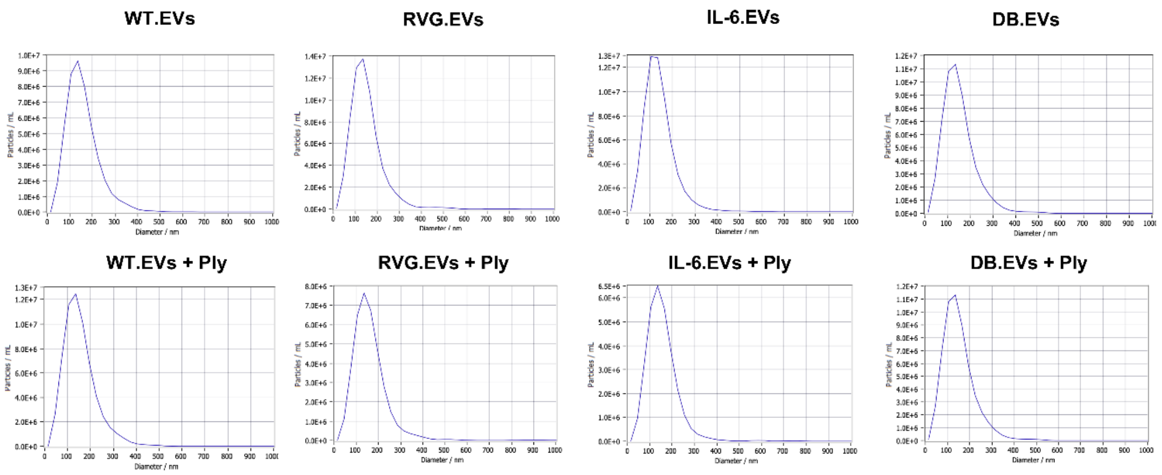


**Supplementary Figure 3. NTA from Ply-sequestering experiments.** Interaction between Ply and the EVs were assessed through size exclusion chromatography (qEV). The eluate was collected and assessed with WB (Figure 3A, B) and NTA was performed on the sample, the ***upper panel*** depicts the size of the EVs without Ply, and the ***lower panel*** depicts the size of the EVs with Ply.

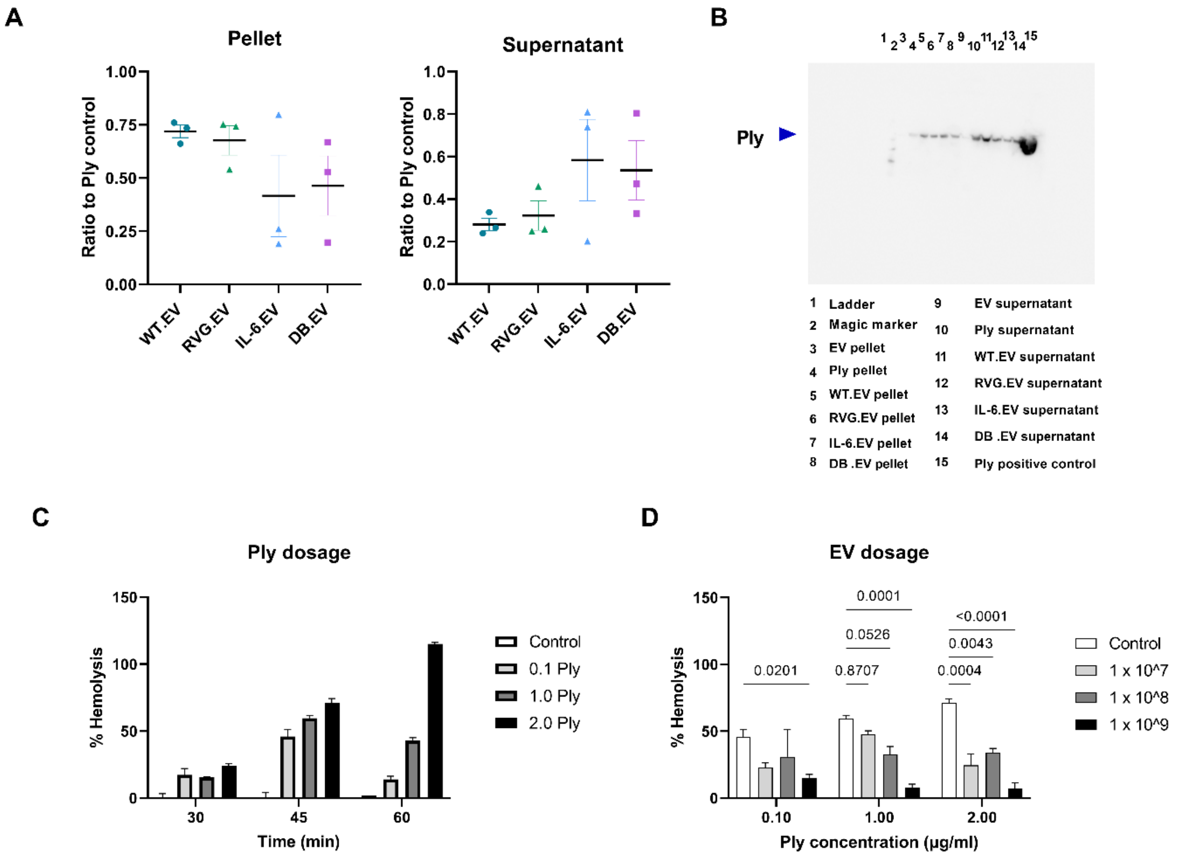


**Supplementary Figure 4. Sequestration of Ply by EVs and titration of Ply and EV dosage in hemolytic assay.** The Ply concentration in the pelleted and supernatant fractions from ultracentrifugation of EV-Ply mixture was assessed by WB (**A, B**). Dose-titration of recombinant Ply concentration to induce hemolysis (**C**). Three different WT.EV concentrations were used to assess their sequestering of Ply (**D**). Experiments were performed in triplicates. Bars depict mean ± SD.

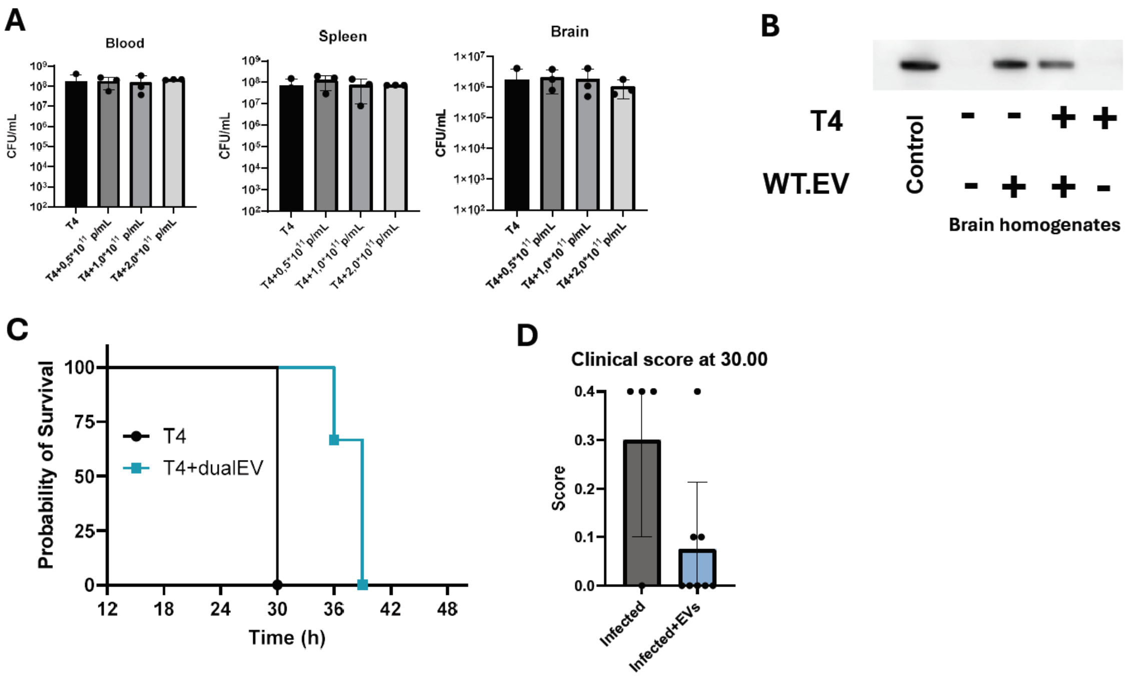


**Supplementary Figure 5. Pilot studies.** Animals were infected with *S. pneumoniae* I.V. then one hour later received EV treatment I.V. Dose titration of the EVs showed that there were no significant differences between bacterial load in the periphery and brain between animal groups (**A**). From brain homogenates we performed western blot analysis for the human CD81 protein. showing that the generated EVs were only in the brain of EV-treated animals and present at 8hrs post-infection (**B**). Assessment of the effectiveness and safety of the DB.EV showed that survival was significantly increased in the DB.EV treated group compared to T4 animal group (**C**). with the majority of DB.EV showed absence or mild symptoms at 30hrs while T4 animal group depicted severe symptoms (**D**). Bars depict mean ± SD.

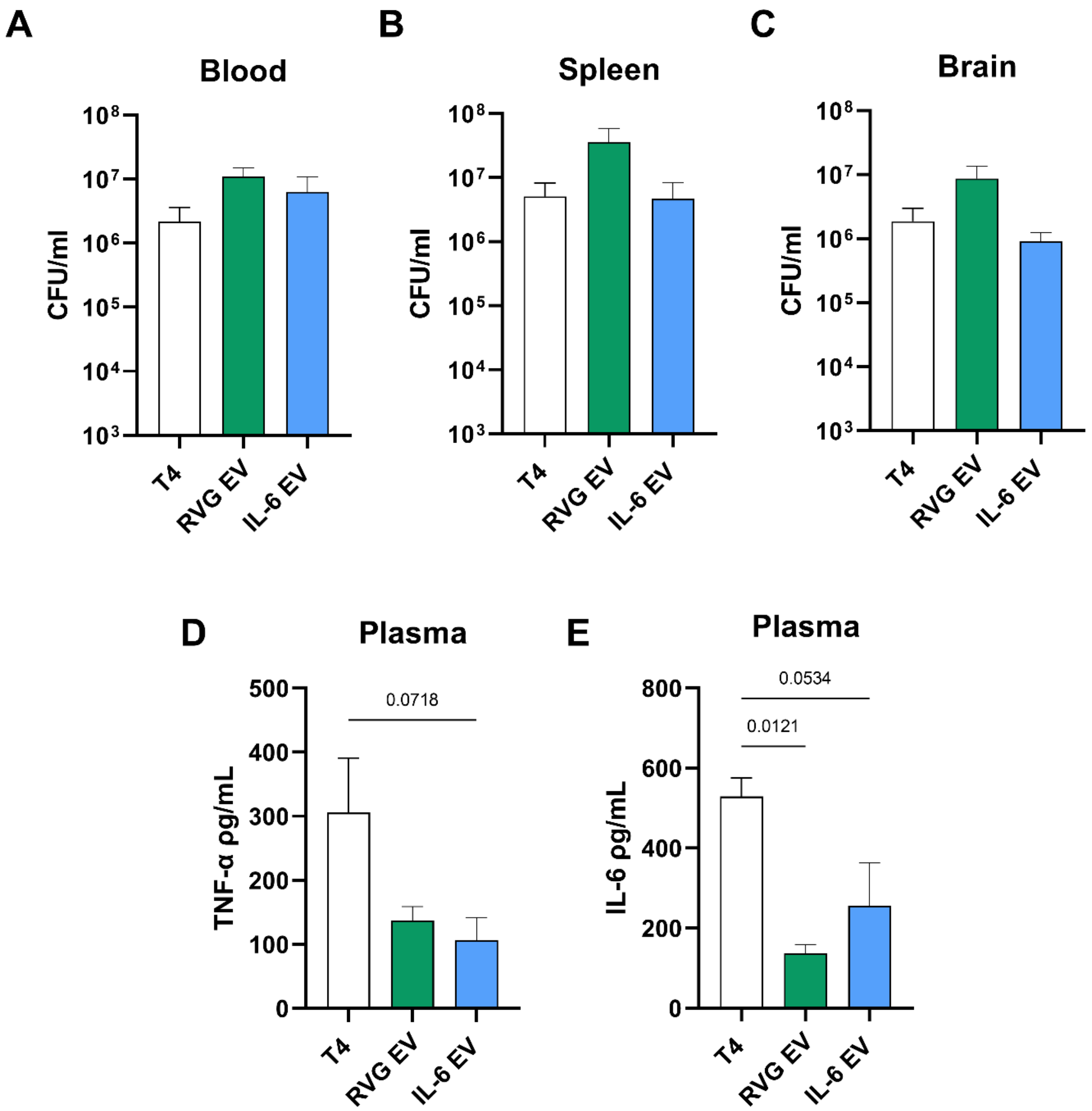


**Supplementary Figure 6. *In vivo* study measuring cytokine levels in plasma.** Animals were infected with *S. pneumoniae* I.V. then one hour later received EV treatment I.V. CFU quantification showed that there was an increase in bacterial load in the blood (**A**), spleen (**B**) and brain (**C**) in RVG.EV treated animals compared to T4 control and IL-6.EV. Despite this both RVG.EV treatment and IL-6.EV treatment reduced plasma levels of TNF-α (**D**) and IL-6 (**E**). Bars depict mean ± SD.

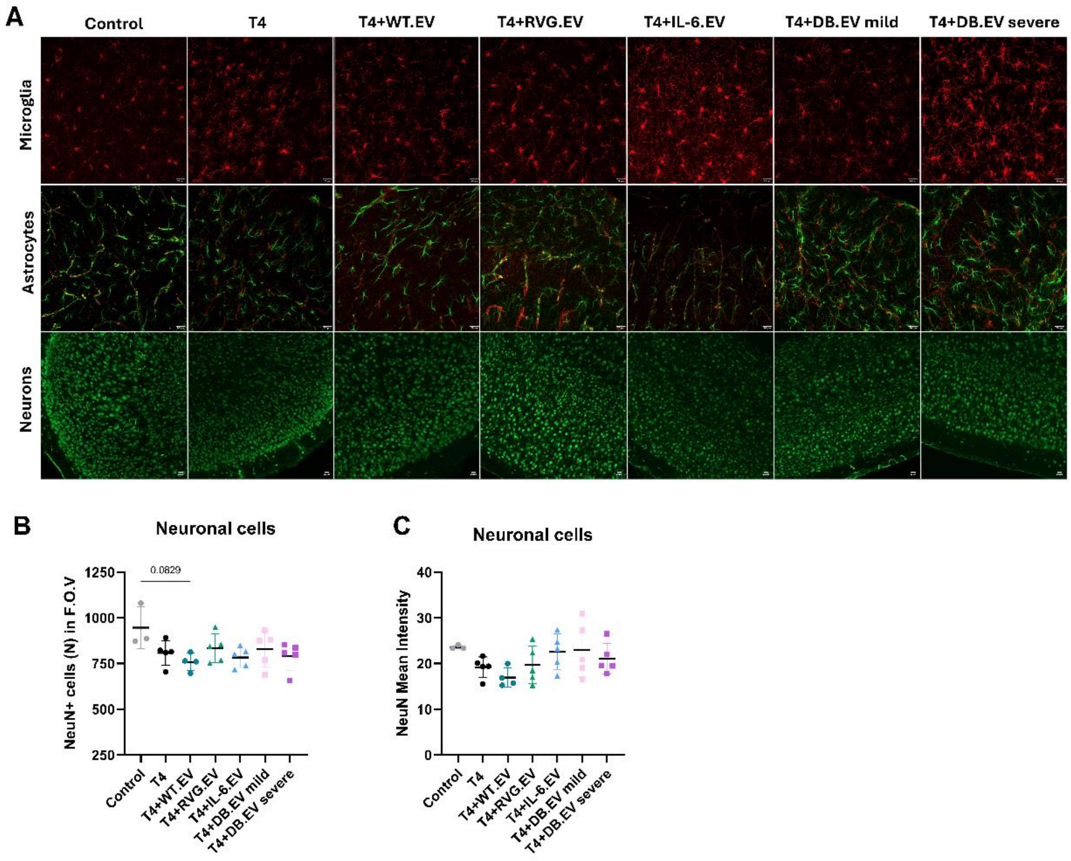


**Supplementary Figure 7. Assessment of neuropathology in pneumococcal meningitis.** Representative images from immunofluorescence analysis (**A**) of microglial cell population (Iba1, red) ***upper panel,*** astrocytes (GFAP, green) and blood vessels (lectin, red) ***middle panel,*** and neurons (NeuN, green) ***lower panel***. No significant difference was observed for the total number of neurons in the F.O.V (**B**) nor intensity of staining (**C**) between animal groups, although WT.EV showed a slight trend in reduction. n= 3 for control, n=4 for WT.EV, and n=5 for RVG.EV, IL-6.EV and DB.EV animal groups. Data were analyzed using a one-way ANOVA and Bonferroni correction for multiple comparisons. Bars depict mean ± SD. Scalebar indicate 20µm.
